# Gel-assisted mass spectrometry imaging

**DOI:** 10.1101/2023.06.02.543480

**Authors:** Yat Ho Chan, Koralege C. Pathmasiri, Dominick Pierre-Jacques, Stephanie M. Cologna, Ruixuan Gao

## Abstract

Compatible with label-free detection and quantification, mass spectrometry imaging (MSI) is a powerful tool for spatial investigation of biomolecules in intact specimens. Yet, the spatial resolution of MSI is limited by the method’s physical and instrumental constraints, which often preclude it from single-cell and subcellular applications. By taking advantage of the reversible interaction of analytes with superabsorbent hydrogels, we developed a sample preparation and imaging workflow named Gel-Assisted Mass Spectrometry Imaging (GAMSI) to overcome these limits. With GAMSI, the spatial resolution of lipid and protein MALDI-MSI can be enhanced severalfold without changing the existing mass spectrometry hardware and analysis pipeline. This approach will further enhance the accessibility to (sub)cellular-scale MALDI-MSI-based spatial omics.

## Main Text

With recent advances in soft ionization techniques, such as matrix-assisted laser desorption/ionization (MALDI) and desorption electrospray ionization (DESI), mass spectrometry (MS)-based spatial analysis (mass spectrometry imaging or MSI) has drastically pushed the technological boundaries of spatial biology research (*1*–*3*). Owing to its compatibility with label-free detection and quantification, MSI has distinct advantages over other imaging modalities in spatial molecular profiling. For example, (sub)cellular-resolution MSI has great utilities in uncovering the molecular constituents behind cellular functions and dysfunctions in intact biological specimens (*4*). In particular, (sub)cellular MALDI-MSI studies can be used to identify endogenous biomolecules and markers associated with metabolic homeostasis, membrane potential regulation, and protein misfolding/aggregation within their original tissue microenvironments.

To date, the spatial resolution of MSI has been limited by the method’s physical and technical constraints. For example, the spatial resolution of MALDI-MSI is limited by the matrix crystal size, which is typically around a few to 10 micrometers (*5, 6*), as well as the instrument’s operating pixel size, which generally falls between 5 to 10s of micrometers, as dictated by the laser spot size and stage precision. A handful of advanced MSI techniques, including secondary ion mass spectrometry (SIMS) (*7*–*9*), custom-built MALDI-TOF mass spectrometers (*10, 11*), and MALDI-2-enhanced sample imaging (*12*–*14*), have demonstrated (sub)cellular-level spatial analysis of intact specimens. However, such methods are often not for daily usage in existing biomedical labs due to the techniques’ limited mass range (*m/z*) and/or availability. These fundamental and practical limitations have slowed down MSI’s application to single-cell and subcellular spatial omics studies.

To allow routine implementation of (sub)cellular MSI in existing labs, here we present a sample preparation and imaging method that overcomes both the physical and instrumental spatial resolution limit of mass spectrometer without modifying its hardware or data analysis pipeline. We name this method <u>G</u>el-<u>A</u>ssisted <u>MSI</u> (or GAMSI), as biological specimens are polymerized (and expanded) using a superabsorbent hydrogel to assist and enhance the MSI capability (**Fig. 1**). Using GAMSI, we show that the spatial resolution of conventional MALDI-TOF mass spectrometers can be enhanced ∼3-4-fold.

**Fig. 1.**
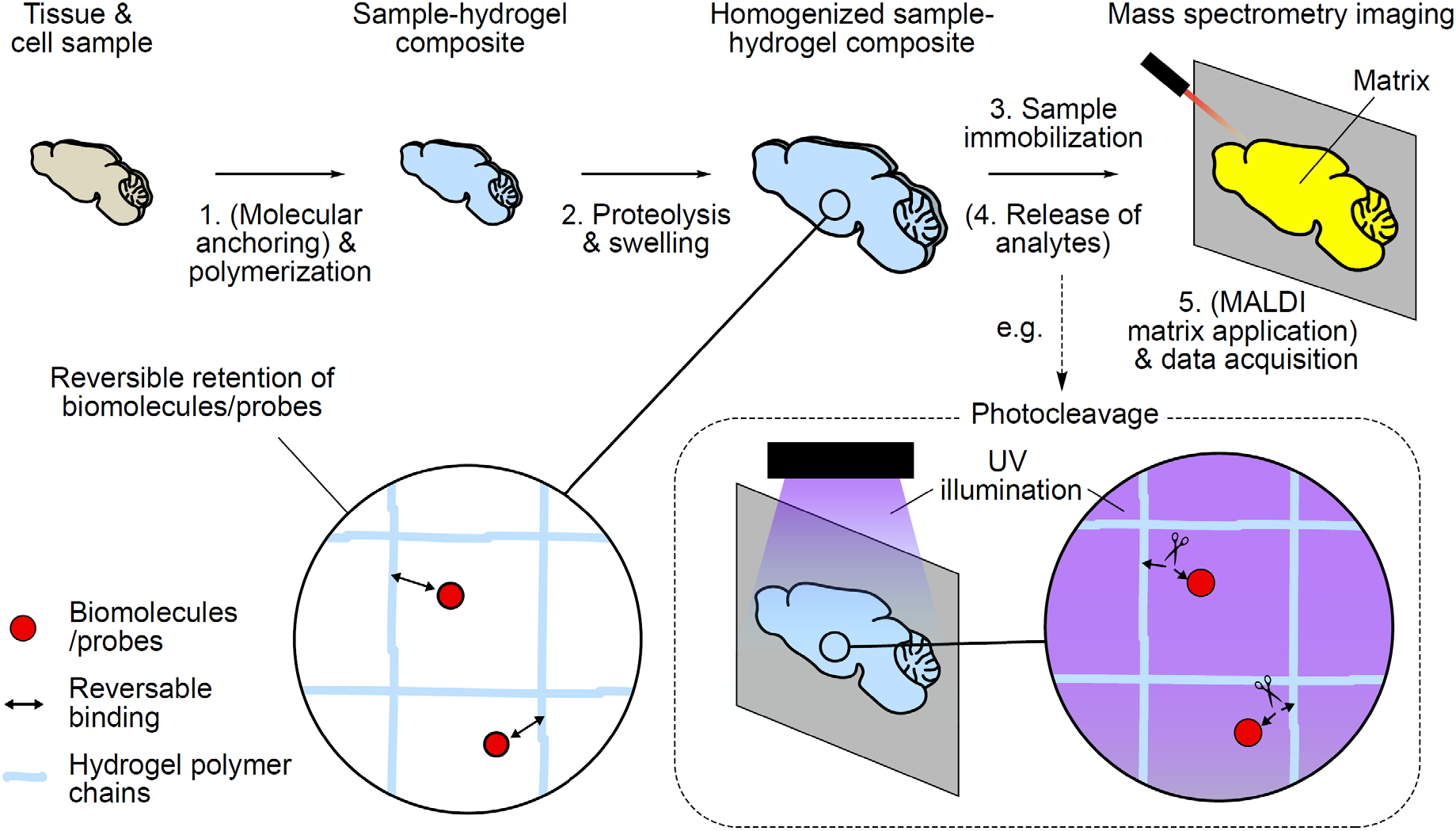
Gel-assisted mass spectrometry imaging (GAMSI). Schematics showing the principle and general workflow of GAMSI. Samples are (1) (optionally) treated with cleavable small-molecule linkers and polymerized to form a superabsorbent hydrogel composite, (2) proteolytically homogenized and expanded, (3) immobilized onto a sample plate, (4) (optionally) subject to cleavage of the small-molecule linkers to release the analytes via, for example, a photocleavage event (dotted box), and (5) (optionally) coated with the MALDI matrix and imaged on a mass spectrometer. The magnified schematics (solid circles) show the reversible tethering/release of biomolecules and probes to/from the hydrogel polymer chains in the sample-hydrogel composite.

### Gel-assisted mass spectrometry imaging (GAMSI)

Pioneering studies combining MSI and mechanical stretching of biological specimens have demonstrated single-cell-sized sample fragmentation and more than an order of magnitude increase in the pixel density for tissue imaging (*15, 16*). More recent studies have shown that the proteomic profile of gel-expanded tissue samples can be determined using existing LC-MS instruments (*17, 18*). These findings have paved the way for our conceptualization of direct MSI on hydrogel-embedded cells and tissues. For GAMSI, we introduce a molecular anchoring strategy that reversibly retains/releases the analysts to/from the hydrogel polymer network (**Fig. 1**). Unlike the covalent protein anchoring scheme adopted by conventional expansion methods, this approach allows *in situ* desorption and detection of biomolecules or probes in the subsequent MSI measurement. We note that the reversible binding of the analytes can take place either physically or chemically. For example, the target biomolecules can be retained via physical interaction with the hydrogel itself or with other molecules covalently anchored to the hydrogel polymer chains. These retained biomolecules can then be released/desorbed and ionized using MALDI. Similarly, the target biomolecules can be chemically tethered to the hydrogel polymer chains using a cleavable small molecule linker. The retained biomolecules can be subsequently untethered by cleaving the linker via, for example, a photocleavage or chemical cleavage event.

### Sample preparation for GAMSI

To demonstrate the concept and working principle of GAMSI, we confirmed that endogenous lipids could be retained and imaged after gel-assisted sample expansion. To enhance lipid retention, we first optimized GAMSI’s sample preparation workflow, including the fixation and proteolysis steps. Fresh-frozen is one of the most commonly used sample formats for MALDI-MSI. For GAMSI, however, we found that a light chemical fixation with paraformaldehyde (PFA) or PFA/glutaraldehyde (GA) was a better choice, because it enhanced the sample integrity for the subsequent polymerization step while maintaining a strong lipid profile comparable to that of the fresh-frozen tissue sample (**fig. S1**) (*19, 20*). Hence, unless otherwise noted, all the GAMSI samples were pre-processed using this light fixation step.

Sample homogenization is a critical step to chemically and/or physically break down the structural integrity of the gelled specimen so that it swells more isotopically in the subsequent expansion step. In the original expansion protocol, proteinase K (proK)-based proteolysis and surfactants were used to homogenize the sample-hydrogel composite (*21*). However, during this process, most (if not all) endogenous lipids are lost due to the strong digestion and the high surfactant concentration. To circumvent this issue, we reasoned that trypsin-based proteolysis with reduced or no surfactants should be sufficient to homogenize thin biological samples (∼5-30 µm) commonly used for MALDI-MSI studies. To test this hypothesis, we treated up to ∼40 µm thick mouse brain slices with a trypsin digestion buffer prepared without any surfactants (**fig. S2**). After homogenizing and expanding the tissue, we observed no visible sample breaking or tearing, which indicated that the sample homogenization was complete and sufficient. To further evaluate the effectiveness of this homogenization step, we characterized the global expansion isotropy of mouse brain slices (∼25 µm) treated with the same trypsin digestion buffer (**fig. S3**). We found that the average expansion error of ∼1-4% of the trypsin-digested tissue slices was comparable to the previously reported ∼1-5% benchmark set by the proK-digested tissue slices (*21, 22*). Combined, these results show that trypsin-based proteolysis is sufficient to homogenize thin tissue samples for the subsequent expansion and imaging. We note that an additional benefit of switching from proK to trypsin digestion is that the latter would provide a more predictable proteolytic profile, which will be critical for untargeted proteomic analysis.

To estimate the percentage of endogenous lipids retained after the gel-assisted expansion, we fluorescently labeled fresh-frozen (as control) and GAMSI-processed mouse cerebellum slices with a phospholipid dye, and measured the fluorescence intensity per normalized unit tissue area (**Fig. 2A, fig. S4**). We found that ∼84% of the phospholipid labeling was preserved in the white matter of mouse cerebellum after the gel-assisted expansion. In addition, ∼82% and ∼100% of the phospholipid labeling was preserved in the molecular layer and the granular cell layer, respectively. These results show that, regardless of the tissue region and lipid abundance, a majority of endogenous lipids were preserved after sample homogenization and expansion.

**Fig. 2.**
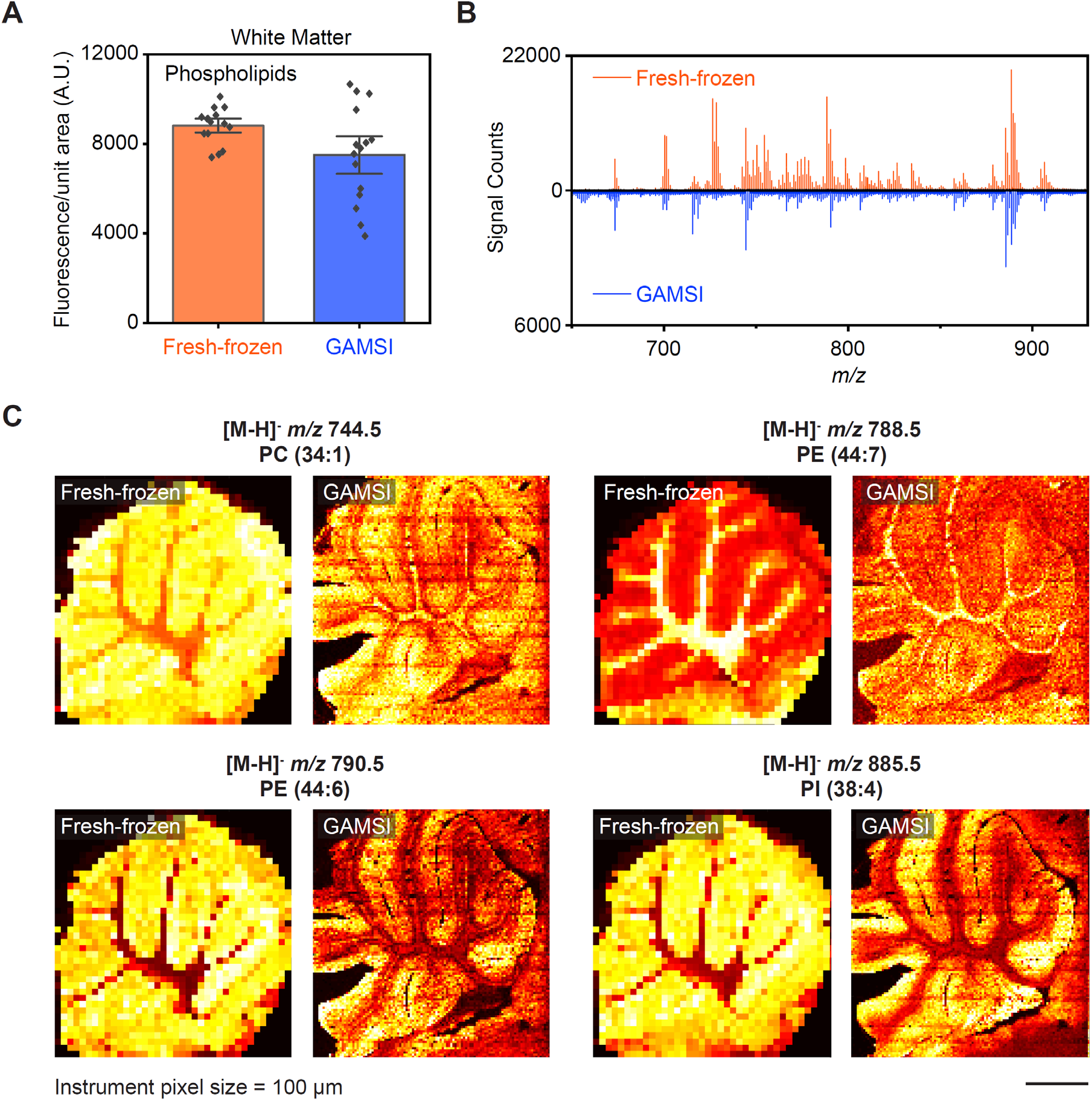
Lipid GAMSI. (**A**) Fluorescence intensity per unit tissue area of fresh-frozen and GAMSI-processed mouse cerebellum white matter, fluorescently labeled using a phospholipid dye [bar height, mean; black dots, individual data points; error bar, standard error of the mean (SEM); n = 15 regions of interest (ROIs) from three brain slices from one animal]. The unit tissue area was normalized to the pre-expansion scale. (**B**) Averaged mass spectra (*m/z* 650-925) of fresh-frozen and GAMSI-processed (PFA/GA-fixed) mouse cerebellum. (**C**) Spatial distributions of representative lipid peaks from the fresh-frozen (left) and GAMSI-processed (PFA/GA-fixed) (right) mouse cerebellum. The instrument pixel size (raster distance) (AB SCIEX 4800) was set at 100 µm. Scale bar: 1 mm (3 mm). Here and after, unless otherwise noted, scale bars are provided at the pre-expansion scale (with the corresponding post-expansion size indicated in brackets). PC: phosphatidylcholine; PE: phosphatidylethanolamine; PI: phosphatidylinositol.

Lastly, we obtained the averaged mass spectra for both the fresh-frozen and GAMSI-processed mouse cerebellum slices on an Applied Biosystems SCIEX 4800 MALDI TOF/TOF Analyzer (“AB SCIEX 4800”) (**Fig. 2B**). We found that the GAMSI-processed sample replicated nearly all the major lipid peaks (in the 650-925 *m/z* range in negative ion mode) of the fresh-frozen sample.

### Enhancing spatial resolution of lipid MSI

Next, we performed lipid GAMSI on mouse cerebellum and ran the same imaging on a fresh-frozen sample as control (**Fig. 2C**). As expected, we observed an enhancement of imaging resolution that corresponded to the ∼3-fold sample expansion. In addition, the fresh-frozen and GAMSI samples showed similar spatial distributions for the major and abundant lipid peaks (**fig. S5-S7**). It is important to note that the observed enhancement of spatial resolution corresponded to nearly an order of magnitude (∼9x = ∼3 x 3) increase in the pixel density, as the number of pixels per normalized unit tissue area scales with the square of the expansion factor. In addition, to validate the identity of the detected lipids, we performed tandem MS on a range of lipid peaks from the GAMSI sample (**fig. S8**). We confirmed that the fragmentation patterns of representative lipid peaks matched those predicted from their molecular structures (*23*–*25*).

These results show that GAMSI can preserve and replicate native molecular and spatial information of fresh-frozen samples.

In principle, the workflow of GAMSI is applicable to all MSI pipelines. Therefore, an increase of spatial resolution by GAMSI should be observed using any MALDI-TOF mass spectrometers.

To validate this, we performed lipid GAMSI on a Bruker rapifleX MALDI Tissuetyper (“Bruker rapifleX”). Similar to the AB Sciex 4800 results, expanded mouse cerebellum images largely captured the native molecular and spatial information of the fresh-frozen counterpart (**fig. S9**-**S10**). Comparably, the ∼4-fold enhancement of imaging resolution corresponded to the observed expansion factor of the sample. These results show that in principle GAMSI can increase the spatial resolution of MALDI-MSI independent of the mass spectrometer hardware.

### Enhancing spatial resolution of lipid-protein MSI

The capability to identify and spatially map multiple types of biomolecules makes MSI a powerful tool for spatial multiomics studies. We next tested if GAMSI could enhance the spatial resolution of multiplexed MSI. As proof-of-concept, we introduced an additional protein labeling step after the lipid GAMSI workflow using an antibody-conjugated, photocleavable mass-tag (*26*). We chose targeted protein imaging for our initial demonstration because the molecular specificity and spatial distribution of antibody labeling has been validated by other methods such as western blot and immunohistochemistry.

**Fig. 3** shows the imaging results of three targeted proteins and a number of endogenous lipids in the expanded mouse cerebellum. The spatial distributions of myeline basic protein (MBP), NeuN, and Synapsin I (SYN-I) matched those validated by immunofluorescence and conventional MSI (*26*). We note that this lipid-protein GAMSI workflow can be scaled up to 10 or more protein targets, because the mass-tag labeling is not limited by spectral overlaps that often hinder molecular multiplexing for fluorescence probes. Furthermore, lipid-protein GAMSI can be extended to untargeted protein imaging, given the recent success in combining mass-tag and untargeted proteomic analyses (*27*).

**Fig. 3.**
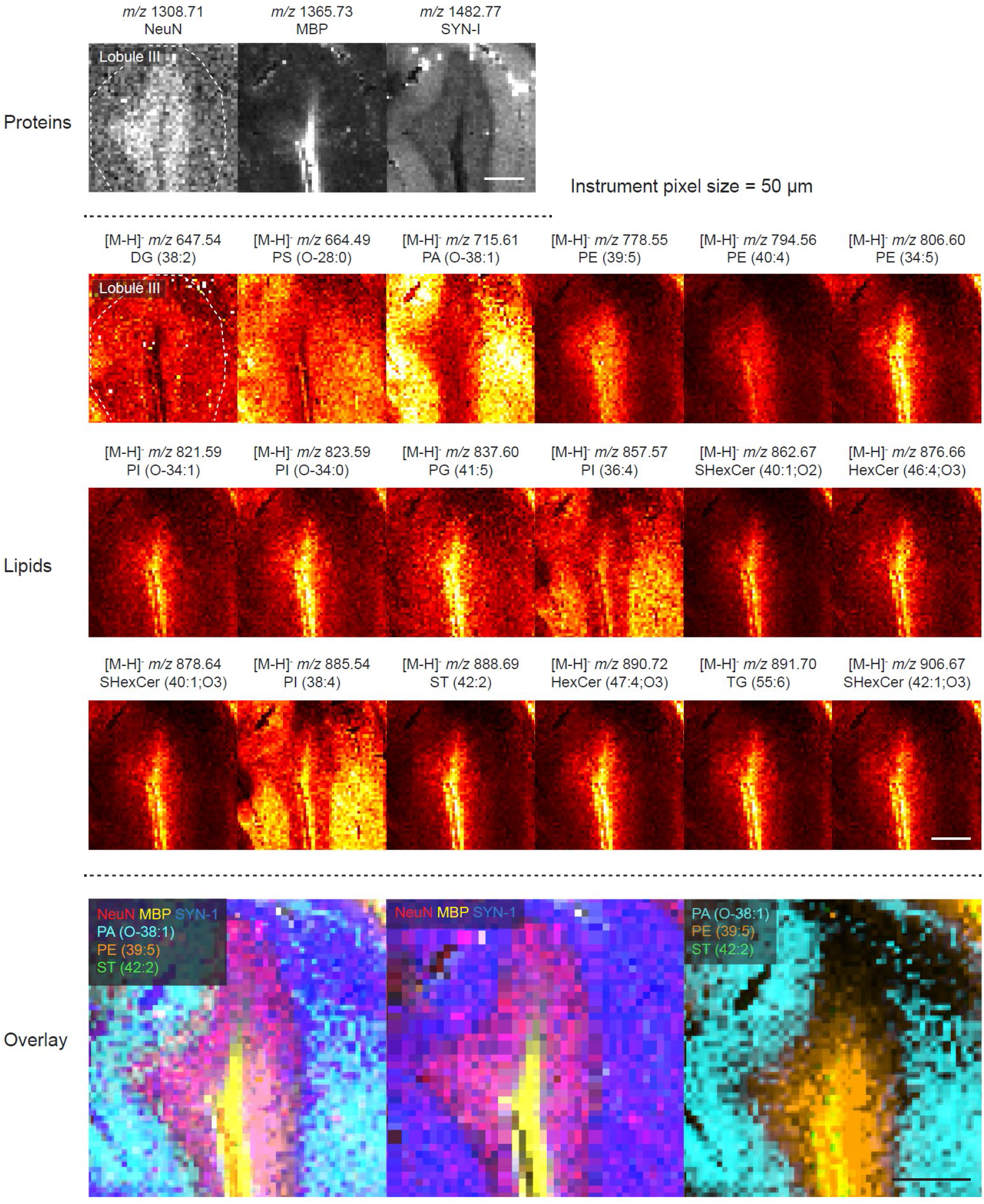
Lipid-protein GAMSI. Spatial distributions of target protein peaks and major lipid peaks from expanded mouse cerebellum (PFA-fixed). The instrument pixel size (Bruker rapifleX) was set at 50 µm. Scale bars: 200 µm (760 µm). DG: diacylglycerol; PS: phosphatidylserine; PA: phosphatidic acids; PE: phosphatidylethanolamine; PG: phosphatidylglycerol; PI: phosphatidylinositol; SHexCer: sulfatides; HexCer: hexosylceramides; ST: sterols; TG: triglyceride.

## Discussion

Currently, the spatial resolution of MALDI-MSI is limited by the physical and instrumental constraints of the method itself, and is often too coarse for single-cell and subcellular studies. In this work, we establish a sample preparation and imaging workflow named GAMSI to overcome these limitations by taking advantage of the analytes’ reversible interaction with a superabsorbent hydrogel. Our results show the spatial resolution of MALDI-MSI can be enhanced by gel-assisted sample expansion. Our study further suggests that such an approach can be applied to multiplexed imaging of different types of molecular targets (e.g., lipids and proteins). This work has strong indication for everyday use of MSI for (sub)cellular-resolution spatial profiling of biomolecules, as its implementation does not require modification of existing MSI hardware or analysis pipeline. Dilution of analytes and dilation of imaging time is the major trade-off for the enhanced spatial resolution. However, as the sensitivity and imaging speed of mass spectrometers continue to improve with the ongoing innovation in the field, even higher enhancement of spatial resolution may be achieved using this approach.

## Acknowledgments

We thank F. Tobias and the Northwestern University IMSERC facility for assistance with mass spectrometry imaging. We thank Chicago Biomedical Consortium (CBC) for access to core mass spectrometry facilities. We thank S. Kwon for assistance with scientific visualization and illustration.

## Funding

This study was supported by

Searle Scholars Program (R.G.)

McKnight Technological Innovations in Neuroscience Award (R.G.)

National Institutes of Health grant UG3MH126864 (R.G.)

University of Illinois Chicago Startup Fund (R.G.)

National Institutes of Health grants R01NS114413 and R01NS124784 (S.M.C.)

National Science Foundation CAREER Award 2143920 (S.M.C.)

## Author contributions

Conceptualization: R.G. and S.M.C.

Methodology: Y.H.C. and R.G.

Investigation: Y.H.C., K.C.P., D.P.-J., and R.G.

Formal analysis: Y.H.C.

Software: Y.H.C.

Visualization: Y.H.C. and R.G.

Funding acquisition: R.G. and S.M.C.

Project administration: R.G.

Supervision: R.G. and S.M.C.

Writing – original draft: R.G. and Y.H.C.

Writing – review & editing: Y.H.C., K.C.P., D.P.-J., S.M.C., and R.G.

## Data and materials availability

All data are available in the main text or the supplementary materials.

## Supplementary Materials

Materials and Methods

Figs. S1 to S10

References(*1*–*32*)

