## Supplementary Material for "Gel-assisted mass spectrometry imaging"

Yat Ho Chan *et al.*

**The PDF file includes:**

Materials and Methods
Figs. S1 to S10
References and Notes

### Materials and Methods

#### 1. Chemicals and Reagents

All reagents were used as supplied unless otherwise noted. Purified water was obtained from a Milli-Q IQ 7000 Ultrapure Water System (Millipore Sigma). All chemicals and reagents were obtained from Millipore Sigma unless otherwise noted.

#### 2. Preparation of Biological Samples

##### *a. Mouse*

All procedures involving mice were performed in accordance with the US National Institutes of Health Guide for the Care and Use of Laboratory Animals and approved by the University of Illinois Chicago Animal Care Committee. Adults heterozygous BALB/cNctr-*Npc1*$^{m1N}$/J mice (Jackson Laboratory) were maintained in a breeding colony at the University of Illinois Chicago. At 7 weeks of age, the mice were euthanized via $CO_2$ asphyxiation followed by decapitation. Whole brains were dissected and immediately frozen on dry ice to maintain tissue integrity. The brains were stored at -80 °C until sectioning.

Brain slices of 25 μm were collected on a CryoStar NX50 Cryostat (Epredia) and thaw-mounted on positively charged microscope slides (MIDSCI) for GAMSI sample preparation, or on stainless steel MALDI plates (Applied Biosystems SCIEX) or ITO-coated glass slides (Hudson Surface Technology, Inc.) for the fresh-frozen control. The sectioned mouse brain slices were stored at -80 °C until further analysis and procedure.

#### 3. Sample Gelation, Digestion, Immunostaining, Expansion, and Immobilization (GAMSI sample preparation)

##### *a. Gelation*

The 25 μm mouse brain slices were thawed to room temperature and fixed with 4% paraformaldehyde (PFA) or 3% PFA/0.1% glutaraldehyde (GA) (for lipid imaging only) at room temperature for 10 min. The fixatives were removed, and the brain slices were washed with 1x PBS three times for 5 min each time. Next, the brain slices were incubated in an Acryloyl-X, SE (AcX, Thermo Fisher) solution (0.1 mg/mL in 1x PBS) at room temperature for 6 hours, and washed with 1x PBS twice for 15 min each time. Gelation of the brain slices was performed following the previous described procedure (*22*, *28*). Briefly, the brain slices were incubated with the monomer solution [1x PBS, 2 M NaCl, 8.625% (w/v) sodium acrylate, 2.5% (w/v) acrylamide, 0.15% (w/v) N,N′-methylenebisacrylamide, 0.01% (w/v) of 4-hydroxy-2,2,6,6-tetramethylpiperidin-1-oxyl (4HT), 0.2% (w/v) of ammonium persulfate (APS), and 0.2% (w/v) of tetramethylethylenediamine (TEMED)] at 4 °C for 30 min and gelled in a humidified 37 °C incubator for 2 hours.

*b. Homogenization*

The gelled brain slices were homogenized in a trypsin (Thermo Fisher) digestion buffer [0.025-0.25% (w/v) in 1x PBS] at 37 °C for 2 days.

*c. Expansion and sample immobilization*

The gelled and digested brain slices were placed in an excess volume of 0.5x PBS for 20 min and then in purified water three times for 20 min each time until the samples were fully expanded. Next, the expanded samples were transferred onto the stainless steel MALDI plates (Applied Biosystems SCIEX) or ITO-coated glass slides (Hudson Surface Technology, Inc.) using a 36 x 60 mm no. 1.5 cover glass (Ted Pella). Finally, the samples were dried under vacuum in a Pyrex vacuum desiccator (Millipore Sigma) filled with Drierite (Millipore Sigma) at room temperature for 4 hours to overnight, and then in a CentriVap Benchtop Vacuum Concentrators (Labconco) at 37 °C for 2 minutes to ensure complete removal of moisture.

*d. Photocleavable mass-tag (PC-MT) modification*

Post-lipid-imaging samples were washed with -80ºC acetone twice for 3 min each time to remove the matrix and then stained with photocleavable mass-tags (PC-MT) (AmberGen, Inc.) on-plate following the protocol provided by the vendor. Briefly, the samples were re-hydrated in a washing sequence of 95% ethanol (EtOH) for 3 min, 70% EtOH for 3 min, and 50% EtOH for 3 min, followed by 1x PBS wash for 10 min. Antigen retrieval was performed using 1x citrate buffer (pH = 6.0) (diluted from 10x citrate buffer, Sigma Aldrich) in a hot water bath for 1 hour at 95ºC, followed by cooling for 30 min at room temperature. The samples were then washed with 1x PBS for 10 min, and incubated with 5 mL of blocking buffer [2% (v/v) normal rabbit serum, 5% (w/v) bovine serum albumin in 1x PBS] for 1 hour at room temperature. Next, the samples were incubated with the PC-MT solution (2.5 µg/ml of each PC-MT in the blocking buffer) overnight at room temperature in a humidified chamber. PC-MTs for NeuN (AP1001201, *m/z* 1308.71), myelin basic protein (MBP) (AP1001200, *m/z* 1365.73), and Synapsin I (SYN-I) (AP1001204, *m/z*

1482.77) were used. After PC-MT incubation, the samples were washed with 1x PBS three times for 5 min each time, then washed with 50 mM ammonium bicarbonate three time for 2 min each time, and dried in a Pyrex vacuum desiccator for 2 hours at room temperature. Finally, the PC-MTs were photocleaved in a UV (365 nm) chamber (AmberGen, Inc.) for 15 min before matrix application.

#### 4. Mass Spectrometry

##### *a. Matrix application*

For lipid imaging on the 4800 Plus MALDI TOF/TOF Analyzer (Applied Biosystems SCIEX), 1,5-diaminonaphthalene (DAN) was applied onto the immobilized and dried sample (on the MALDI plate) via sublimation using a previously described homemade sublimation apparatus (*29*, *30*). Briefly, 50 mg of DAN was dissolved in 2 mL of acetone, aspirated onto the bottom of the sublimation flask, and blow-dried using nitrogen to form a thin layer. Next, a hotplate, to which the sublimation flask was placed, was set to 105 °C while a digital thermometer was placed in contact with the bottom of the flask to monitor the temperature. An ice slush was added to the cold finger of the apparatus, to which the MALDI plate was adhered on the underside with copper tape. Finally, the sublimation apparatus was placed under vacuum at 80 mTorr using a rough pump, and the DAN matrix was sublimed for 2 min. The amount of matrix deposited on the sample was determined as mass per centimeter square.

For lipid imaging on the rapifleX MALDI Tissuetyper (Bruker), 9-aminoacridine (9AA) solution was sprayed to the sample with a HTM M3+ sprayer (HTX Technologies LLC) using the following settings: Nozzle temperature = 60ºC; nozzle velocity = 1200 mm/min; flow rate = 0.12 mL/min; 9AA concentration = 10 mg/mL; number of passes = 8; track spacing = 2 mm; nitrogen gas pressure = 10 psi. 9AA solution was prepared by dissolving 100 mg of 9AA to 10 mL of solution of 75% ACN, 25% purified water, followed by filtering.

For PC-MT imaging on the rapifleX MALDI Tissuetyper (Bruker), α-cyano-4-hydroxycinnamic acid (CHCA) solution [7 mg/mL in 50% ACN, 50% purified water, and 0.1% trifluoroacetic acid (TFA)] was sprayed to the sample with the HTM M3+ sprayer using the following settings: Nozzle temperature = 79ºC; nozzle velocity = 1200 mm/min; flow rate = 0.1 mL/min; CHCA concentration = 7 mg/mL; number of passes = 10; track spacing = 2 mm; nitrogen gas pressure = 10 psi.

*b. Mass spectrometry imaging*

Initial lipid imaging was performed with a 4800 Plus MALDI TOF/TOF Analyzer equipped with a 200 Hz Nd:YAG pulse laser (355 nm) (Applied Biosystems SCIEX). The instrument was externally calibrated and operated in negative ion reflection mode to acquire data between mass ranges (*m/z*) of 500-1000 using DAN as matrix. The number of laser shots per pixel was set at 50 and the raster distance between each pixel was set to 100 µm using the 4800 Imaging Tool v.3.2 (https://ms-imaging.org/wp/4000-series-imaging).

Additional lipid imaging was performed with a rapifleX MALDI Tissuetyper equipped with a Smartbeam 3D 10 kHz Nd:YAG (355 nm) laser (Bruker). The instrument was operated in negative ion reflection mode to acquire data between mass ranges (*m/z*) of 500-2000 using 9AA as matrix. The number of laser shots per pixel was set at 200 and the raster distance between each pixel was set to 50 µm using FlexImaging (Bruker).

For PC-MT imaging, the rapifleX MALDI Tissuetyper was operated in positive ion reflection mode to acquire data between mass ranges (*m/z*) of 1000-2000 using CHCA as matrix. The number of laser shots per pixel was set at 200 and the raster distance between each pixel was set to 50 µm using FlexImaging.

Data processing including region-of-interest determination, average mass spectrum extraction, and image generation was conducted using MSiReader (ver. 1.02) (*31*). All images presented were normalized using Total Ions Count (TIC). Lipid assignments were made by comparing mass measurements to the LIPID MAPS database (www.lipidsmaps.com), with an allowed mass tolerance of *m/z* = +/- 0.05. Image registration across lipid and protein images were performed using TrackEM2 (ver. 1.0a 2012-07-04) plugin on ImageJ distribution Fiji (ver. 1.53t).

*c. Tandem mass spectrometry (MS/MS)*

MS/MS was performed with the 4800 Plus MALDI TOF/TOF Analyzer (Applied Biosystems SCIEX) in negative ion reflection mode using DAN as matrix. The instrument was operated in the 2 kV operation mode to allow unimolecular decay.

#### 5. Fluorescence Microscopy

##### *a. Expansion isotropy*

To evaluate the spatial isotropy of sample homogenization and expansion, 25 μm mouse brain slices were thawed, fixed with 3% PFA/0.1% GA, and washed with 1x PBS three times for 5 min each time following the standard GAMSI sample preparation workflow described earlier. The fixed brain slices were incubated in a detergent-free blocking buffer [5% (v/v) normal goat serum (NGS) in 1x PBS] at room temperature for 6 hours. Next, the brain slices were incubated in the primary antibody (rabbit anti-NF-200, N4142-.2ML, Millipore Sigma) solution (1:100 dilution with the detergent-free blocking buffer) overnight at room temperature. The brain slices were then washed with the blocking buffer four times for 30 min each time, and incubated in the secondary antibody (goat Alexa Fluor 568-conjugated anti-rabbit antibody, A11011, Thermo Fisher) solution (1:200 dilution with the detergent-free blocking buffer) for 2 days at room temperature. Finally, the brain slices were washed with 1x PBS (or the detergent-free blocking buffer) four times for 30 min each time and stored in 1x PBS.

Pre-expansion fluorescence images were obtained using a Nikon spinning disk confocal system (CSU-W1, Yokogawa) with a 40x 1.15 NA water immersion objective (Nikon). After pre-expansion imaging, the brain slices were gelled, homogenized, and expanded following the standard GAMSI sample preparation workflow described earlier. The expanded brain slices were then imaged using the same confocal microscope system and objective across regions of interest (ROIs) corresponding to those obtained from the pre-expansion imaging. Lastly, pre- and post-expansion image registration, vector deformation field generation, and root mean square (r.m.s) error calculation of all point-to-point measurements were performed using a custom MATLAB code as previously described (*32*).

##### *b. Lipid retention*

For fresh-frozen (non-expanded) samples, 25 μm mouse brain slices were thawed to room temperature and immediately stained with HCS LipidTOX™ Red Phospholipidosis Detection Reagent (Thermo Fisher) for phospholipid labeling (1:200 dilution with 1x PBS) overnight at room temperature. The brain slices were then fixed with 3% PFA/0.1% GA at room temperature for 2 hours to maintain tissue integrity. Finally, the fixatives were removed, and the brain slices were washed with PBS three times for 20 min each time.

For expanded samples, 25 μm mouse brain slices were thawed, fixed, gelled, homogenized following the standard GAMSI sample preparation worked. The samples were then incubated with

HCS LipidTOX™ Red Phospholipidosis Detection Reagent (1:200 dilution with 1x PBS) overnight at room temperature expanded with purified water three times for 20 min each time.

Wide-field epifluorescence images of both the fresh-frozen and the expanded lipid-stained brain slices were taken with a 10x 0.45 NA air objective (Nikon) with 2 ms exposure. Total pixel values per pre-expansion unit tissue area of the white matter, molecular layer, and granular cell layer of cerebellum were measured to represent fluorescence intensity per unit tissue area.

*c. Comparison of trypsin and proteinase K (proK) digestion*

40 µm mouse brain slices (from one mouse brain, perfused and fixed with 4% PFA) were used to evaluate different digestion conditions for the sample homogenization step. The first brain slice was subject to the standard GAMSI sample preparation. Briefly, the fixed brain slice was incubated in the detergent-free blocking buffer [5% (v/v) normal goat serum (NGS) in 1x PBS] at room temperature for 6 hours. Next, the brain slice was incubated in the primary antibody (rabbit anti-NF-200, N4142-.2ML, Millipore Sigma) solution (1:100 dilution with the detergent-free blocking buffer) overnight at room temperature, and then washed with the blocking buffer four times for 30 min each time. The brain slice was then incubated in the secondary antibody (goat Alexa Fluor 568-conjugated anti-rabbit antibody, A11011, Thermo Fisher) solution (1:200 dilution with the detergent-free blocking buffer) for 2 days at room temperature, washed with 1x PBS (or the blocking buffer) four times for 30 min each time, and stored in 1x PBS. Finally, the brain slice was gelled, homogenized with trypsin digestion buffer, expanded, and imaged following the standard GAMSI sample preparation workflow. Pre- and post-expansion fluorescence images were obtained using a Nikon spinning disk confocal system (CSU-W1, Yokogawa) with a 4x 0.20 NA air objective (Nikon).

The second brain slice was subject to the proExM protocol (*22*, *28*). Briefly, the fixed brain slice was permeabilized with 0.1% (w/v) Triton X-100 in 1x PBS for 15 min and incubated in a blocking buffer [5% (v/v) normal goat serum (NGS) and 0.1% (w/v) Triton X-100 in 1x PBS] at room temperature for 6 hours. Next, the brain slice was incubated in the primary antibody (rabbit anti-NF-200, N4142-.2ML, Millipore Sigma) solution (1:100 dilution with the blocking buffer) overnight at room temperature, and washed with the blocking buffer four times for 30 min each time. The brain slice was then incubated in the secondary antibody (goat Alexa Fluor 568-conjugated anti-rabbit antibody, A11011, Thermo Fisher) solution (1:200 dilution with the blocking buffer) overnight at room temperature, washed with 1x PBS (or the blocking buffer) four times for 30 min each time, and stored in 1x PBS. Finally, the brain slice was gelled, homogenized with proK digestion buffer, expanded, and imaged following the proExM protocol. Pre- and post-expansion images were obtained using the Nikon spinning disk confocal system with a 4x 0.20 NA air objective.

### Supplementary Figures

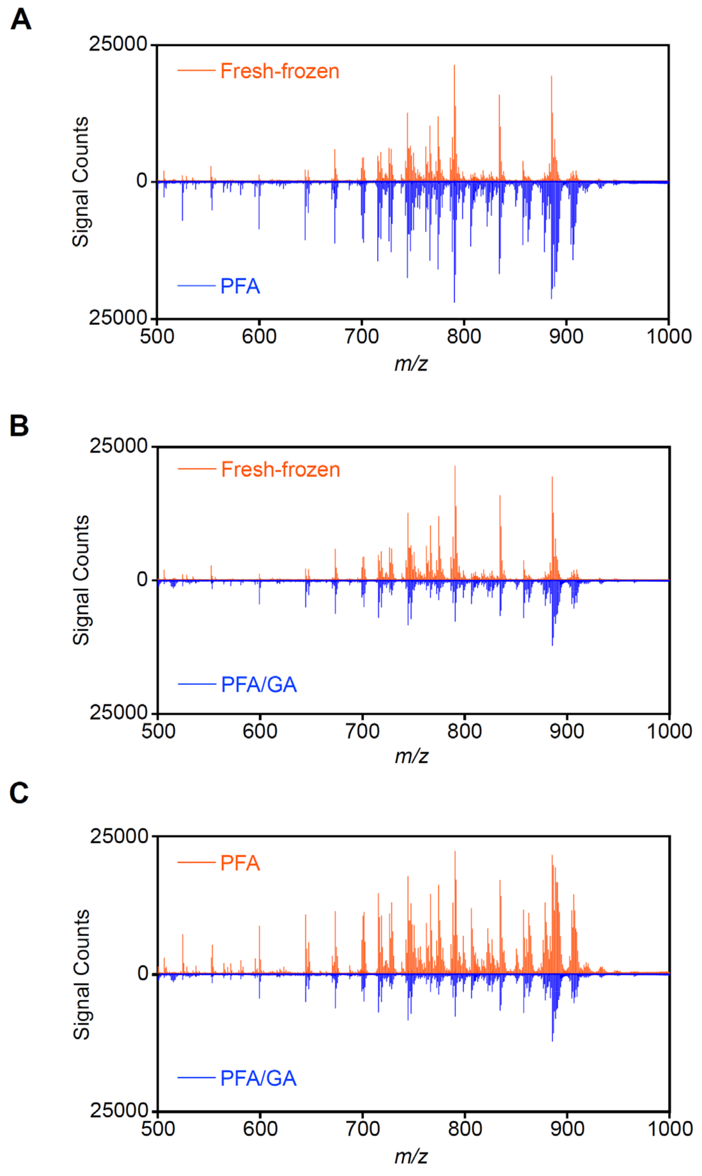


**Fig. S1. Averaged mass spectra (*m/z* 500-1000) of fresh-frozen, PFA-fixed (non-expanded), and PFA/GA-fixed (non-expanded) mouse cerebellum.** Comparisons between (**A**) fresh-frozen and PFA-fixed, (**B**) fresh-frozen and PFA/GA-fixed, and (**C**) PFA and PFA/GA-fixed sample mass spectra are shown. The mass spectra were collected using an Applied Biosystems SCIEX 4800 MALDI TOF/TOF Analyzer ("AB SCIEX 4800"). The instrument pixel size was set at 100 µm.

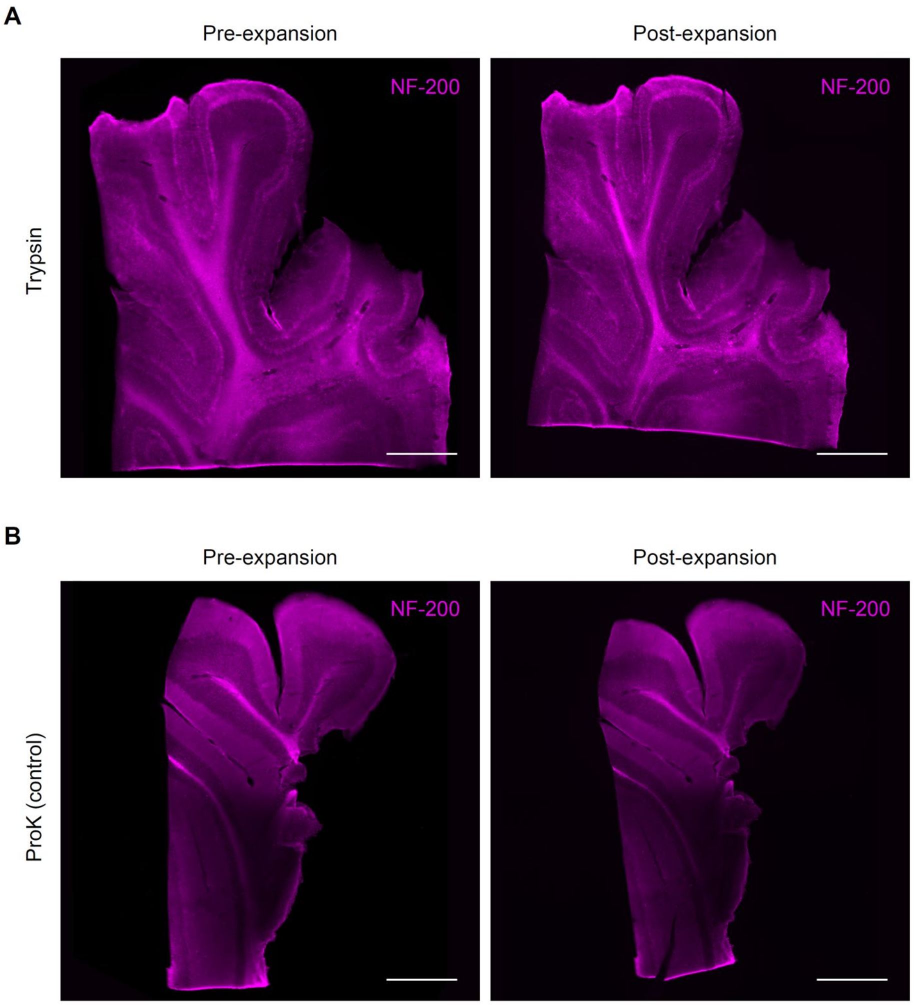


**Fig. S2. Trypsin and proteinase K (proK) digestion of thin tissue slices**. Pre- (left) and post-(right) expansion images of ~40 µm mouse brain slices digested using the (**A**) trypsin (with no surfactants) and (**B**) proK (with surfactants, control) digestion buffers. The brain slices were fluorescently labeled using neurofilament (NF)-200 antibodies. Scale bars, 500 µm (1.9 mm). Here and after, unless otherwise noted, scale bars are provided at the pre-expansion scale (with the corresponding post-expansion size indicated in brackets).

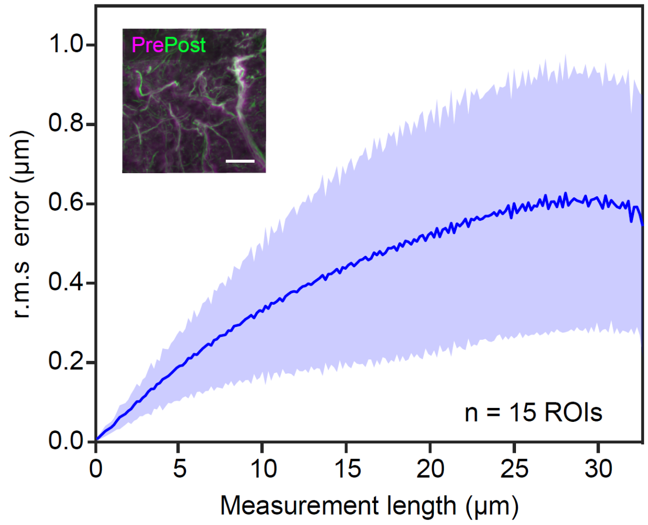


**Fig. S3. Expansion isotropy analysis.** Root-mean-square (r.m.s.) expansion error of ~25 µm mouse brain slices homogenized by trypsin proteolysis [blue line, mean; shaded area, standard deviation; n = 15 regions of interest (ROIs) from three brain slices from two animals]. Inset: Non-rigidly registered and overlaid pre- (magenta) and post-expansion (green) images used for the r.m.s. error analysis. Scale bar, 5 µm (15 µm).

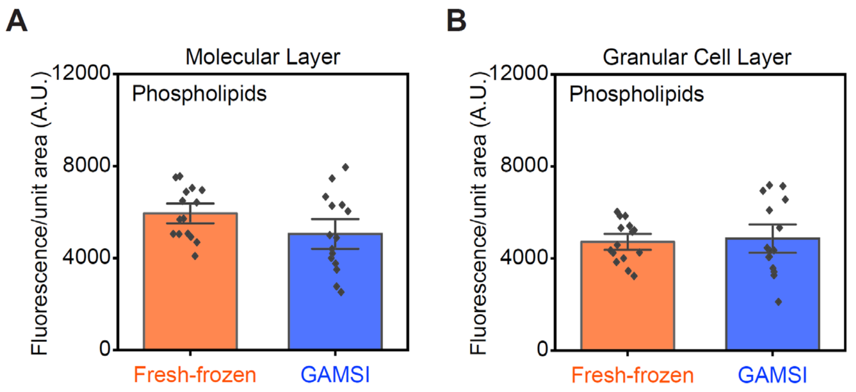


**Fig. S4. Lipid retention in GAMSI-processed mouse brain slices (phospholipid labeling).** Fluorescence intensity per unit tissue area of fresh-frozen and GAMSI-processed mouse cerebellum (**A**) molecular layer and (**B**) granular cell layer, fluorescently labeled using a phospholipid dye [bar height, mean; black dots, individual data points; error bar, standard error of the mean (SEM); n = 15 regions of interest (ROIs) from three brain slices from one animal]. The unit tissue area was normalized to the pre-expansion scale.

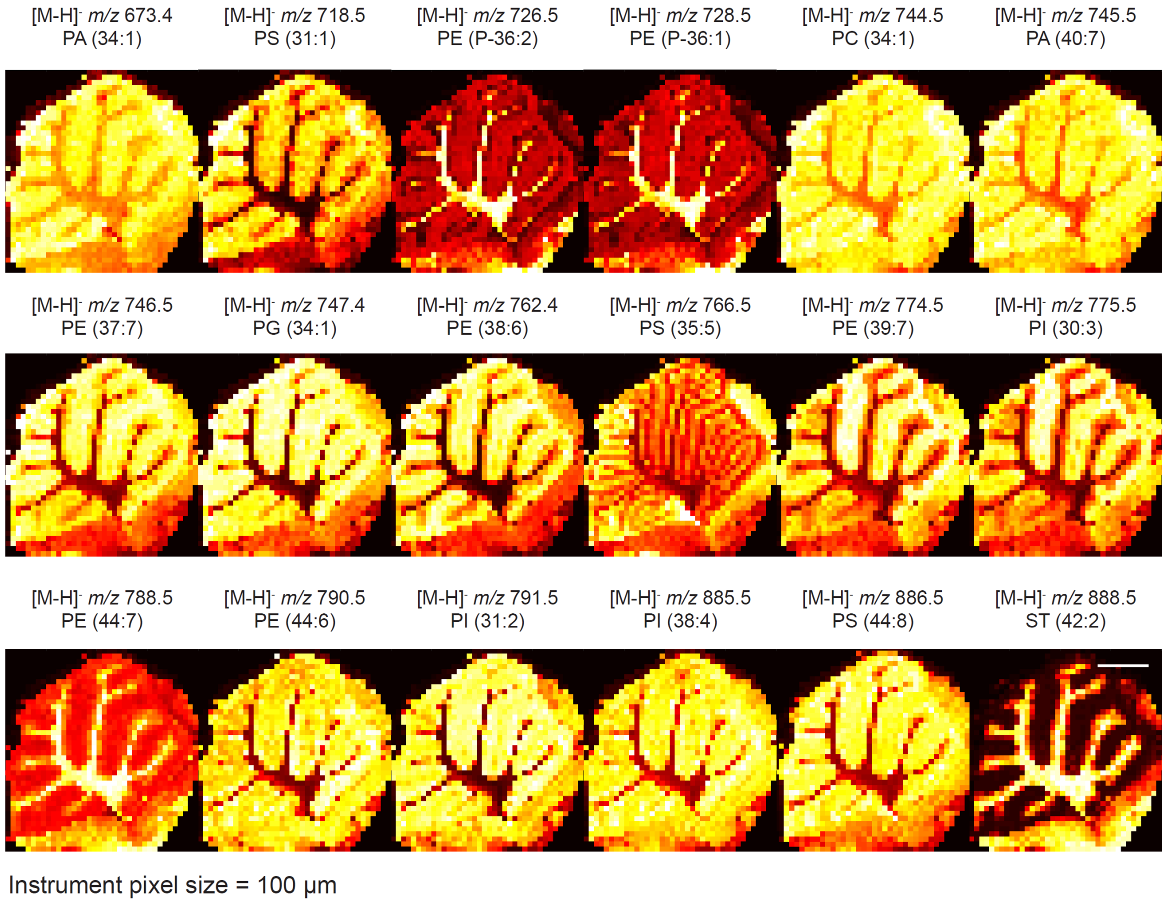


**Fig. S5. Spatial distributions of major lipid peaks from fresh-frozen mouse cerebellum.** The sample was imaged using AB SCIEX 4800 with an instrument pixel size of 100 µm. Scale bar: 1 mm. PA: phosphatidic acids; PS: phosphatidylserine; PE: phosphatidylethanolamine; PG: phosphatidylglycerol; PI: phosphatidylinositol; PC: phosphatidylcholine; ST: sterols.

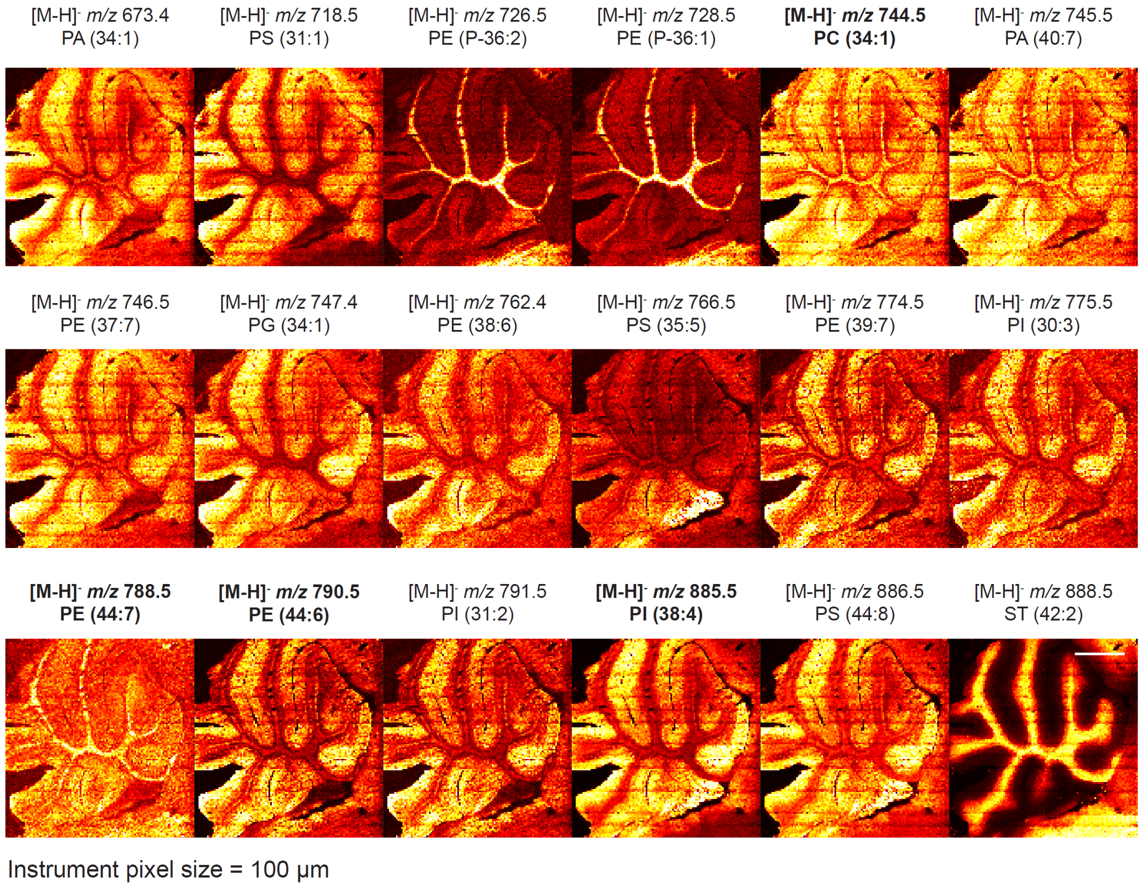


**Fig. S6. Spatial distributions of major lipid peaks from lipid GAMSI of PFA/GA-fixed mouse cerebellum.** The GAMSI-processed sample was imaged using AB SCIEX 4800 with an instrument pixel size of 100 µm. Scale bar: 1 mm (3 mm). Lipid names include both those predicted from the lipid database (LIPID MAPS) and confirmed by MS/MS (bolded). PA: phosphatidic acids; PS: phosphatidylserine; PE: phosphatidylethanolamine; PG: phosphatidylglycerol; PI: phosphatidylinositol; PC: phosphatidylcholine; ST: sterols.

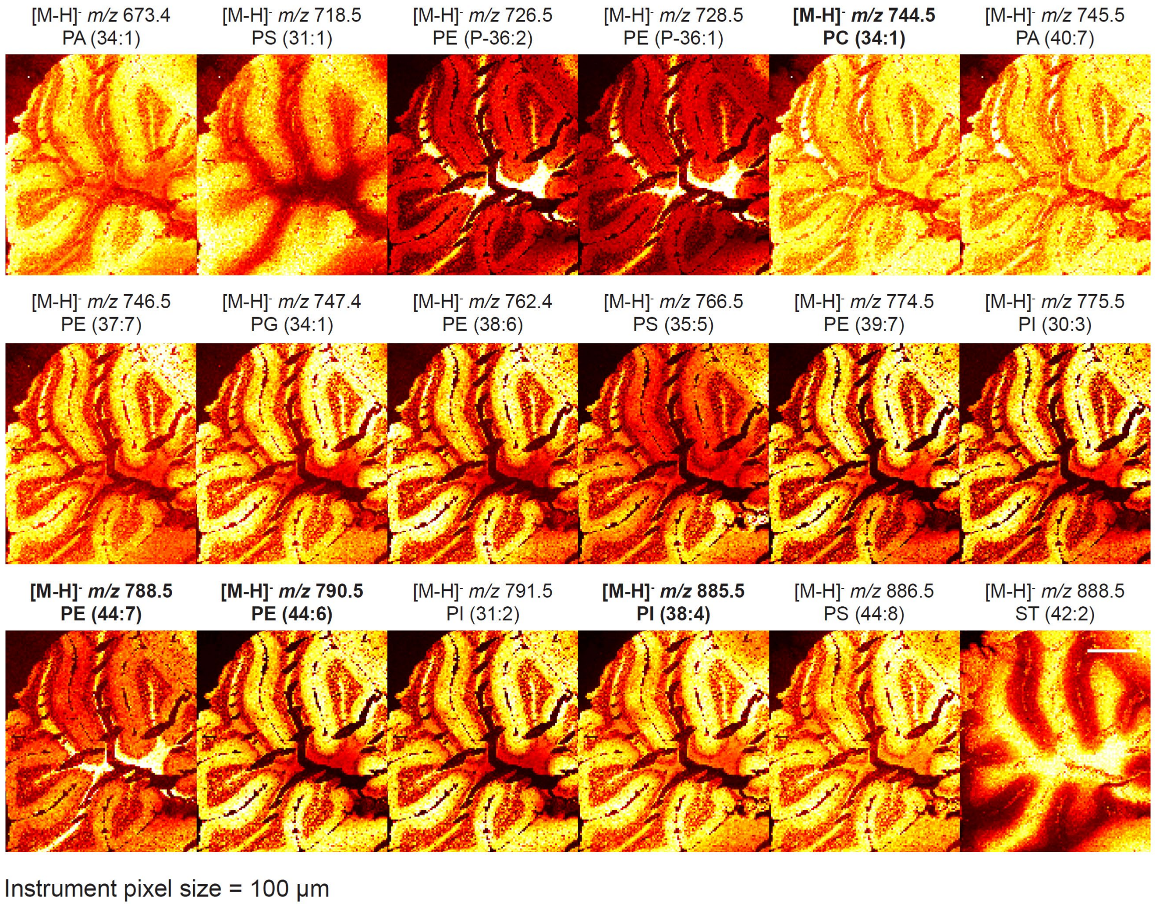


**Fig. S7. Spatial distributions of major lipid peaks from lipid GAMSI of PFA-fixed mouse cerebellum**. The GAMSI-processed sample was imaged using AB SCIEX 4800 with an instrument pixel size of 100 μm. Scale bar: 1 mm (4 mm). Lipid names include both those predicted using the lipid database (LIPID MAPS) and confirmed by MS/MS (bolded). We note that the observed tissue tears were from cryo-sectioning. PA: phosphatidic acids; PS: phosphatidylserine; PE: phosphatidylethanolamine; PG: phosphatidylglycerol; PI: phosphatidylinositol; PC: phosphatidylcholine; ST: sterols.

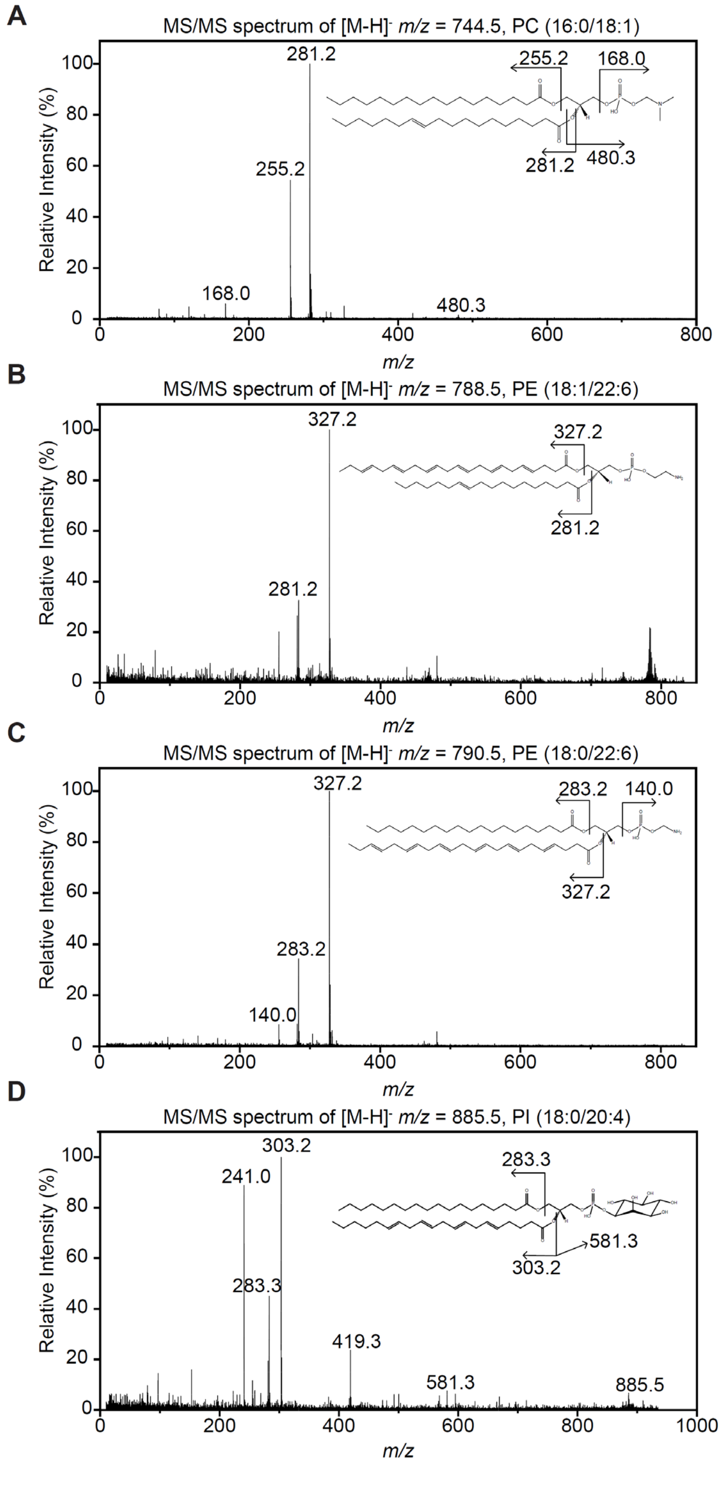


**Fig. S8. MS/MS spectra of representative lipid peaks from GAMSI-processed mouse cerebellum**. MS/MS spectra of *m/z* = (**A**) 744.5, (**B**) 788.5, (**C**) 790.5, and (**D**) 885.5 peaks. The spectra were collected using AB SCIEX 4800 with an instrument pixel size of 100 µm.

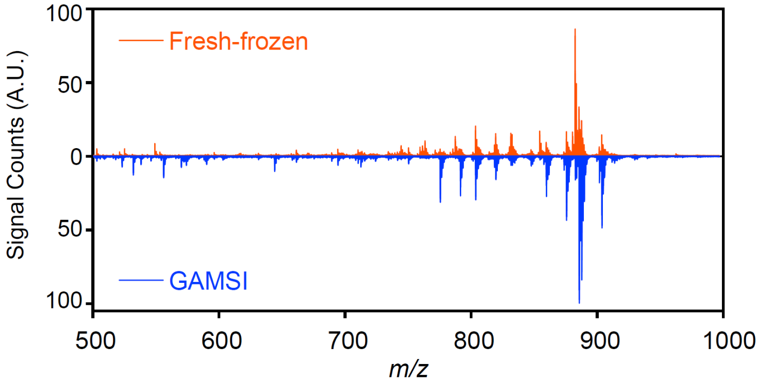


**Fig. 9 Averaged mass spectra (*m/z* 500-1000) of fresh-frozen and GAMSI-processed mouse cerebellum.** The mass spectra were collected using a Bruker rapifleX MALDI Tissuetyper ("Bruker rapifleX"). The instrument pixel size was set at 50 µm.

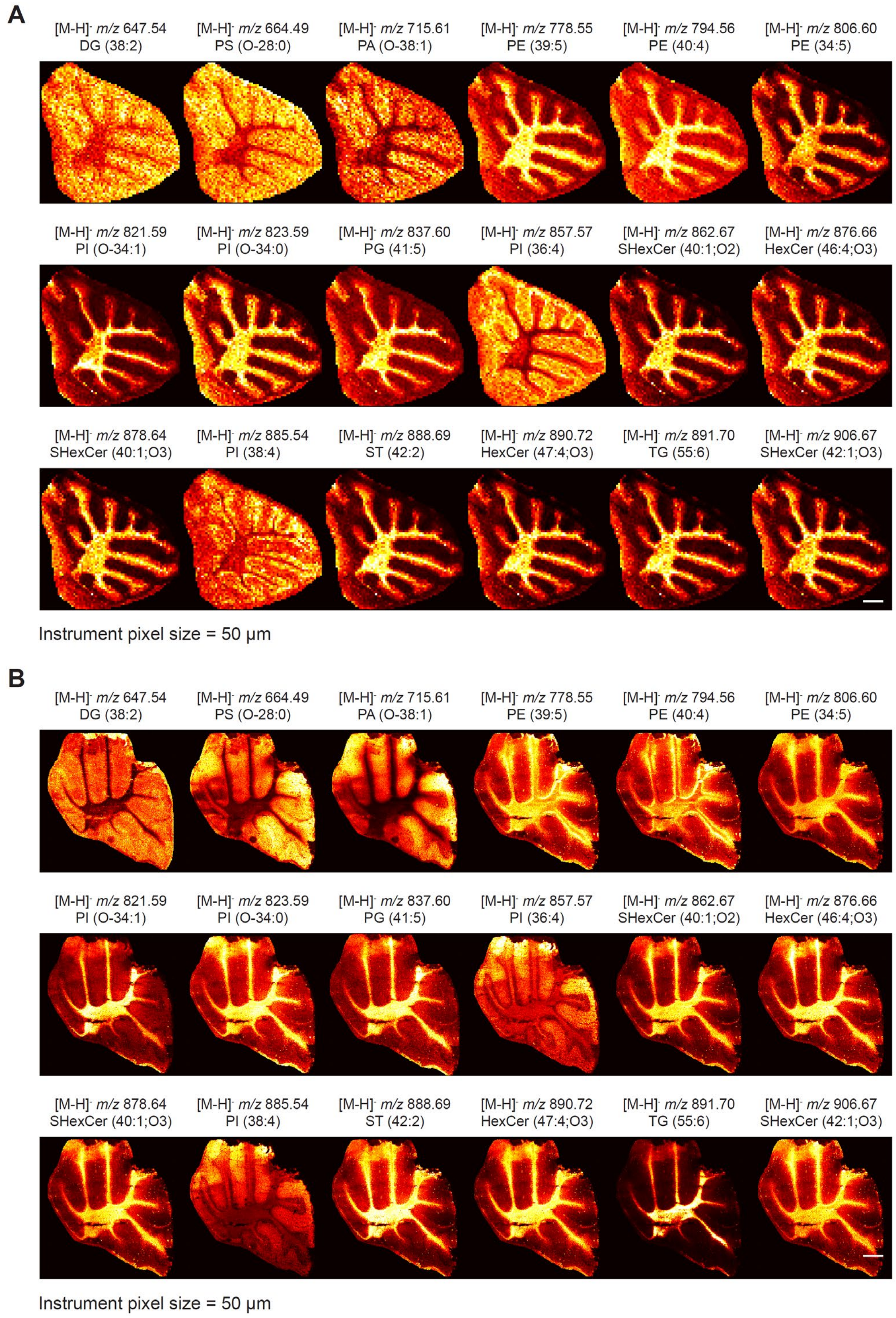


**Fig. S10. Spatial distributions of major lipid peaks from fresh-frozen and GAMSI-processed (PFA-fixed) mouse cerebellum.** Both the (**A**) fresh-frozen and (**B**) GAMSI-processed samples were imaged using Bruker rapifleX with an instrument pixel size of 50 µm. Scale bar: 500 µm (1.9 mm). DG: diacylglycerol; PS: phosphatidylserine; PA: phosphatidic acids; PE: phosphatidylethanolamine; PG: phosphatidylglycerol; PI: phosphatidylinositol; SHexCer: sulfatides; HexCer: hexosylceramides; ST: sterols; TG: triglyceride.
